# Mapping, stacking and characterization of a novel resistance haplotype against *GpaV_vrn_*-virulent *Globodera pallida* derived from wild potato relative *Solanum brevicaule*

**DOI:** 10.64898/2026.10.27.762148

**Authors:** D. Langendoen, J.M. de Boer, I. Mager, H. Room, J-J. Willig, Y. Do, A. N.O. Rookmaker, A. Winiarczyk, R.H. Zwinderman, J.E. Tammes, M.G. Sterken

## Abstract

Potato production is threatened by pests and pathogens. Among the most harmful are the potato cyst nematodes (PCN), including *Globodera pallida.* The major control measures are potato cultivars harbouring resistance genes. However, virulent nematode populations are breaking through the most broadly applied resistance genes, and therefore novel resistance genes are required to keep control of *G. pallida*. In this study we map a resistance locus in wild potato relative *Solanum brevicaule* to the bottom of chromosome 11 (42.83 - 42.85 Mb) using a k-mer based bulked segregant analysis and name the resistant haplotype *GpaXI_brc_.* The presence of the *GpaXI_brc_* marker in the potato genome significantly increases resistance against a *GpaV_vrn_*-virulent *G. pallida* population (AMPOP01). Furthermore, the *GpaXI_brc_* haplotype was stacked with three other resistance haplotypes: *GpaXI_tar_*, *GpaV_vrn_,* and *GpaIX_vrn_*. Stacking of the novel *GpaXI_brc_* haplotype to these other resistance haplotypes resulted in a significant increase in resistance. The largest effect was seen in combinations with “major” resistance haplotypes *GpaV_vrn_* + *GpaIX_vrn_* and *GpaXI_tar_* + *GpaXI_brc_*. To determine whether we are dealing with a novel allele, the resistance mechanism underlying the GpaXI_brc_ haplotype was described after performing an *in vitro* bioassay. The resistance mechanism was different from *GpaXI_spl_* as described in literature, which is also localized at the bottom of chromosome 11. Therefore, we conclude that we found a novel resistance allele originating from *S. brevicaule*.

## Introduction

Potato production is threatened by pests and pathogens. Among the most harmful are the potato cyst nematodes (PCN) *Globodera pallida* and *Globodera rostochiensis* (J. T. Jones et al., 2013). PCN are biotrophic, sedentary endoparasitic nematodes that feed on potato roots. PCN extract nutrients and assimilates that are essential for plant growth and thus symptoms are similar to those associated with nutrient deficiency, such as stunted growth, yellowing of leaves and reduced yields. Heavy infestations are seen in the field as patches that have a reduced growth and show latent flowering (Price et al., 2024). Yield losses due to PCN infestation are estimated to be 9% worldwide (Turner & Rowe, 2006), but locally heavier losses of up to 80% have also been reported (Oerke, 2006). It is therefore important to manage and control PCN. Often applied practices are crop rotation, nematicides, and resistant cultivars (Sikora et al., 2021). Of these, cultivation of cultivars with resistance genes is among the most potent and easily applied control measures against PCN.

In the past decades, many resistance haplotypes have been discovered. The PCN resistance genes that were cloned all belong to the NBS-LRR family (Ernst et al., 2002; Paal et al., 2004; Van Der Vossen et al., 2000). Of the resistance loci, for which we do not know the causal gene, most are located in resistance gene clusters (regions with a higher number of predicted NB-LRRs) within the potato genome (Gebhardt & Valkonen, 2025). For an excellent overview of virtually all known PCN resistance loci, a recent review by Gartner et al. (Gartner et al., 2021) can be consulted. The effect of PCN resistances on nematodes is variable and dependent on the nematode population and genetic background of the potato cultivar (Price et al., 2024). The first described resistance haplotype was H1 against *G. rostochiensis* (Ellenby, 1952). Application of H1 led to the discovery of *G. pallida* and subsequently various resistance haplotypes against *G. pallida* were described (F. G. W. Jones et al., 1970). In particular *GpaV_vrn_* (also referred to as Gpa5), originating from *Solanum vernei,* has been used extensively in breeding programmes and is readily available in cultivars such as Iledher and Seresta (Eoche-Bosy et al., 2017; Rouppe van der Voort et al., 2000; Schaveling, te Molder, et al., 2026). Throughout the years, farmers have mainly relied on this resistance haplotype to protect their potatoes against *G. pallida* (Schaveling, te Molder, et al., 2026). Other resistances to *G. pallida* are rarer in modern potato cultivars, such as *GpaXI_tar_* (Tan et al., 2009) or *GpaIX_vrn_* (Bryan et al., 2002; Rouppe van der Voort et al., 2000).

Nevertheless, the implementation of R gene-based strategies revealed that eradication of the entire population could not be achieved (Trudgill, 1985). Using *GpaV_vrn_* as an example, artificial selection of *G. pallida* over multiple generations on cultivars carrying *GpaV_vrn_* resistance produced virulent populations able to reproduce on the formerly resistant host. (Beniers et al., 2019; Fournet et al., 2013; Turner et al., 2009; Varypatakis et al., 2020).

Furthermore, recent reports of virulence on potato farms in Germany and the Netherlands confirm that natural populations are no longer controlled by this gene (Den Nijs & Van Heese, 2019; Niere et al., 2014; Schaveling, te Molder, et al., 2026). With the commercial introduction of *GpaV_vrn_* in the cultivar Seresta in 1996, inadvertently a next generation of resistances would be needed in the future.

Here we report a new source of resistance against *G. pallida* originating from the wild diploid potato relative *Solanum brevicaule*. We mapped the resistance to a QTL on the right arm of chromosome XI. We used the CoSSA bulked segregant method (Prodhomme et al., 2019) to find a genetic marker, named *GpaXI_brc_*. This genetic marker can be used in potato breeding programs for enhanced genotype selection to simplify – or even eliminate – the need for phenotypic screening on *G. pallida* resistance. Importantly, we show this resistance is functional against recently reported *GpaV_vrn_*-virulent *G. pallida* and thereby can help in the control of the recently emerging field populations.

## Material and methods

### Plant material

Firstly, in 2016, *Solanum brevicaule* clone KA 2014-9087 (CGN18030) was found to be resistant to *GpaV_vrn_*-virulent *Globodera pallida* population AMPOP01 in a pot-test bioassay. Before further characterization, we confirmed the absence in this genotype of the *G. pallida* resistance haplotypes that were reported in literature (Gartner et al., 2021) by PCR analysis (**Table S1**). Genotype KA 2014-9087 was crossed with susceptible diploid *S. tuberosum* KA 2007-4176 (RH 06-976). From here on, various crosses were performed (A pedigree overview of the material used in this research is depicted in **Fig. S1**). The resulting seeds were sown under greenhouse conditions in 2017. Of 60 genotypes, tubers were harvested for resistance testing. This mapping population was named P 2018-3004.

Secondly, resistance pyramiding was investigated using additional crosses. In 2019, one of the most resistant and best-performing genotypes within P 2018-3004, KA 2018-3227, was crossed twice to incorporate additional resistances against *G. pallida*, namely *GpaV_vrn_* (van Eck et al., 2017) and *GpaIX_vrn_* (Bryan et al., 2002) via a cross with KA 2015-5546 and *GpaXI_tar_* (Tan et al., 2009) via a cross with KA 2016-2537. The population segregating for *GpaV_vrn_* and *GpaIX_vrn_* was named P 2020-5501, and the population segregating for *GpaXI_tar_* was named P 2020-2006. Seeds of these crosses were sown in the greenhouse and per genotype, a single tuber was harvested for field propagation in 2021 in Valthermond, The Netherlands. Six tubers per genotype were further propagated under field conditions in 2022. In 2023, three tubers per genotype were used in a resistance test. Genetic markers for *GpaV_vrn_*, *GpaIX_vrn_* and *GpaXI_tar_* based on (Tan et al., 2009), (van Eck et al., 2017) and (Bryan et al., 2002) were used for identification and selection.

Thirdly, two additional clones carrying the *S. brevicaule* resistance were retained for investigations, namely a sibling of the *S. brevicaule* parent from the BSA cross (KA-2014- 9088) and a second resistant clone from the mapping population (KA 2018-3228).

### Sequencing of samples

Based on the pot test results, a resistant bulk (R-bulk; RS<5%; n=18) and a susceptible bulk (S-bulk; RS>15%; n=18) were composed from genotypes of mapping population P 2018- 3004 (n=60). DNA was isolated from freeze dried young leaf material of both bulks using a KingFisher Flex DNA Extraction System (Thermo Fisher Scientific, Bleiswijk, the Netherlands) in combination with a Sbeadex Plant DNA Purification kit (LGC BioSearch Technologies, Berlin, Germany). Illumina sequencing was done by Novogene GmbH (Munich, Germany) and by GenomeScan B.V. (Leiden, The Netherlands), producing 150 bp paired end fastq files.

Similarly, a number of genotypes were sequenced from young leaves: KA 2014-9087 is the susceptible parent, KA 2014-9088 served as a substitute for the original resistant parent (KA 2014-9087), which was lost during breeding, KA 2018-3227 and KA 2018-3228 are individuals from the resistant bulk, KA 2007-4176 is the susceptible parent. The sequencing depths of the samples are shown in **Table S2**.

### PCN populations

In this study, various populations of *G. pallida* were used. The *GpaV_vrn_*-virulent AMPOP populations were sampled from agricultural fields growing partially resistant potato cultivars in the Netherlands around 2015 and were described previously (Schaveling, te Molder, et al., 2026). AMPOP01 has been propagated on susceptible cultivar Desiree since.

AMPOP01S2D was propagated (selected) on partially resistant cultivar Seresta for two generations, after which it has been propagated on Desiree. Both populations showed selection on the *Gp-pat-1* locus in *G. pallida*, conferring virulence on *GpaV_vrn_* (Schaveling, van Rijt, et al., 2026).

### Resistance testing

The pot-test bioassay used to research potato genotypes for resistance to *G. pallida* is based on (Den Nijs, 2024). Tubers were cut into similar sizes using a melon scoop and kept in plastic bags overnight to allow a cork layer to form. Tuber pieces were then planted in 0.7 L pots in 700 g of silver sand/clay/hydrograin mixture (4:0.7:1) supplemented with fertilizer (N:P:K = 12:10:18) and Steiner solution (Steiner, 1961). Directly after planting the tuber cutting, pots were inoculated with *G. pallida* AMPOP01 (five living larvae and eggs/g of soil), obtained by crushing the cysts using a rotating pestle and a glass vial. Plants were watered daily using a sprinkler for the first two weeks. After two weeks, all plants were watered simultaneously using ebb and flood irrigation, biweekly supplemented with fertilizer (Fertigo 6-3-6/Fertigo 2-4-9) from week five onwards. Plants were grown until mature cysts could be observed on susceptible control plants upon uprooting (approximately after 12 weeks). The air-dried contents of the pots were sieved, and cysts were extracted using Fenwick’s method (EPPO, 2013). Cysts were then counted, and the relative susceptibility (RS) was calculated using the following formula:

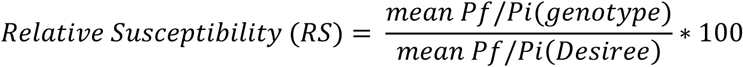

In addition to the pot test, plants were also researched in an *in-vitro* infection assay. Sprouts were removed from the tubers and surface sterilized using 70% ethanol (15 seconds), 2% NaOCl + 150 µL/L Tween20 (10 min) followed by three washing steps in sterilized tap water. Plants were maintained on MS medium (Murashige & Skoog, 1962) supplemented with 30 g/L sugar (MS30) in a climate chamber (18°C, 16h light/8h dark cycle). Nodal segment cuttings of the plants (∼1 cm) were planted on Gamborg B5 (Duchefa) medium containing 1.5% bacto-agar and 2 g/L sucrose. Ten days after planting, the rooting cuttings were inoculated with approximately 120 sterile J2 larvae of *G. pallida* (AMPOP01). Cysts were hatched at room temperature in the dark on a 100 μm sieve in a Petri dish with tap water (one week) followed by tomato root diffusate (another 7 days). J2s were purified by centrifugation on 70% sucrose (2000 rpm, 4 min.), after which the middle layer containing the J2s was sterilized with 1.0% NaN_3_ and 0.8% HgCl_2_. After sterilization, the J2s were kept in sterile tap water until use. Before inoculation, J2s were transferred to a 0.7% (w/v) gelrite solution. Five droplets of 1 µL containing 20 J2s were inoculated near the roots. Droplets of 0.7% (w/v) gelrite solution without nematodes were used as mock control. Male/female formation was assessed at 28 dpi, cysts were counted at 63 dpi.

Since prerequisites for ANOVA were not met for the resistance tests, Kruskal-Wallis Rank Sum tests or PERMANOVA tests were performed to study significant differences between groups. For Kruskal-Wallis Rank Sum test, the kruskal() function within the R-package “agricolae” was used with standard settings (alpha = 0.05, p.adj = “none”). For PERMANOVA the dist() and adonis2() functions were used from the R package “vegan” with method = "euclidean" and permutations = 999.

### Bulked Segregant Analysis

A k-mer based Bulked Segregant Analysis (BSA) was used for the QTL mapping, following the CoSSA principle of Prodhomme et al. (2019). The kmc3 package was used for creating 31 bp k-mer databases from the Illumina sequence samples in Table S2. These databases contained the k-mers having 2 or more copies in the sequence reads sets. Because the ends of the sequence reads do not yield k-mers, the overall k-mer depths of the samples will be less than the read depths, and estimated k-mer depth values are shown in Table S2. The k- mer distributions of the sample databases are shown in Fig. S2. They fit to the estimated k- mer depths of Table S2. The kmc3 software was also used for the k-mer operations on these databases (Kokot et al., 2017).

The standard k-mer database processing approach for a CoSSA is given by the formula:

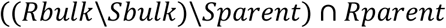

In an initial analysis, we followed this formula, with the modification that sibling KA 2014- 9088 was used as substitute for the lost R parent. However, this procedure did not give a clear result and showed more than one positive chromosome region in the BSA, possibly because this was a sibling and not the actual parent, resulting in false k-mer signals not related to the resistance haplotype (data not shown). Therefore, another alternative for the R- parent was chosen, by using two sequenced clones from the R bulk, KA 2018-3227 and KA 2018-3228, for the intersection step. The complete series of processing steps is depicted in Table S3. In the final cleanup, k-mers were selected that had a depth value range of 10-30 in KA 2018-3227. This interval corresponds with the peak of simplex k-mers in this genotype **(Fig. S2)** and should contain the resistant haplotype.

Contrary to the procedure used by Prodhomme et al. (2019), the localisation of the QTL was performed by mapping reads, instead of k-mers. The database with the 751878 selected k- mers was used by the filter function of kmc tools, to fetch the k-mer positive reads from the reads set of the resistant bulk. This filtering produced a mix of single and paired reads. A subset of 33,724 read pairs then was identified with a custom script and was mapped to the DM 6.1 reference genome (Pham et al., 2020). The number of mapped reads per 10 kbp interval was determined with sambamba software (Tarasov et al., 2015) and the chromosomal distribution of the paired read counts was visualized using R.

To calculate significances based on the k-mer count data, we used an exponential distribution after log_10_ transformation. In this analysis bins with 0 counts assigned to them were excluded. We used the EnvStats package to calculate the rate of the distribution and the pexp function to assign significances to k-mer counts per bin.

### PCR assay development

The reads of the sequenced samples were mapped to the potato DM 6.1 reference genome (Pham et al., 2020) using bwa software with standard settings (Li & Durbin, 2009). Variants were called from the *.bam alignment files with freebayes software (Garrison & Marth, 2016).

For SNP comparison between samples, SNP dosage values for the selected genomic regions were extracted from the *.vcf variant files with a custom Python script.

Graphical genotyping was applied to identify Single Nucleotide Polymorphisms (SNPs) on chromosome 11 between 42.5 Mb and 44.5 Mb, corresponding to the peak found after the BSA. Selected SNPs were specific to the resistant bulk and resistant genotypes KA 2014- 9088, KA 2018-3227 and KA 2018-3228 (**Figs. 2B and S3)**, while being absent in the susceptible bulk and other susceptible genotypes. For the latter selection, a database with SNP profiles of 283 genotypes was used. Seven PACE markers were developed by 3cr Bioscience (UK).

For SNP genotyping analysis, PCR Allele Competitive Extension (PACE) Genotyping 2x Master Mix (Low ROX – 25 nM) (3cr) was used, following the manufacturers protocol with some minor adaptations. The assay mix was prepared by combining two allele-specific primers and one common primer at a concentration of 100 µM in a ratio of 1:1:2, respectively. Primer sequences were mentioned in **Table S4**. The genotyping mix was then loaded into 384-well PCR plates, with 2.5 µl of master mix, and 0.07 µl of assay mix per well. Subsequently, 2.5 µl of DNA was added to each well.

Thermal cycling was performed using SensoQuest Labcyclers with 384-well thermoblocks, following this protocol: an initial denaturation step at 94°C for 15 minutes, followed by ten cycles of denaturation at 94°C for 20 seconds and annealing with a ramped temperature starting from 61°C to 55°C, decreasing by 0.6°C per cycle, for 60 seconds. This was followed by 26 cycles of denaturation at 94°C for 20 seconds and annealing at 55°C for 60 seconds. Finally, 9 cycles of denaturation at 94°C for 20 seconds and annealing at 57°C for 60 seconds were performed. PCR products were assessed by endpoint fluorescence using a QuantStudio 6 Flex Real-Time PCR System (Thermo Fisher Scientific).

Single-marker regression analysis was conducted in R to evaluate the association between each marker and relative susceptibility. A linear model was fitted using the lm() function, with relative susceptibility as the response variable and the marker as the predictor. The coefficient of determination (R²) was calculated as:

*R*² = 1 − (*SSresidual*/ *SStotal*).

To adjust for model complexity, the adjusted R² was computed as:

*Adjusted R*² = 1 − [(1 − *R*²)(*n* − 1) / (*n* − *p* − 1)],

where *n* is the number of observations and *p* is the number of predictors.

## Results

### A *S. brevicaule* derived resistance locus on chromosome 11

To find new sources of resistance, in 2016, a pot test screening was performed using *GpaV_vrn_*-virulent *G. pallida* population AMPOP01. Two *S. brevicaule* clones (KA 2014-9087 and KA 2014-9088, originating from CGN18030, **Fig. 1A**), scored an average replication ratio (P_f_/P_i_) of 0.71 and a relative susceptibility (RS) of 1.54% when compared to Desiree. In 2017, a follow-up pot test was performed to confirm the resistance. This test was inoculated with the slightly more virulent population AMPOP01S2D (Schaveling, van Rijt, et al., 2026) (**Fig. 1B**). This resulted in a P_f_/P_i_ of 1.78 and an RS of 3.59% when compared to Desiree. Based on these consecutive tests we concluded that CGN18030 contains resistance against *G. pallida*.

After identifying a potential resistance gene donor, we set out to further characterize the resistance and determine whether a resistance-associated locus could be mapped. To that end, KA 2014-9087 was crossed with an *S. tuberosum* clone, KA 2007-4176. of which the pedigree does not contain known sources of resistance. The resulting mapping population (P 2018-3004) of 60 different genotypes was tested for *G. pallida* resistance in a pot test using AMPOP01 and subsequent genetic analysis by bulked segregant analysis.

We determined resistance in P 2018-3004 and found average RS scores ranged from 0 to 33.8% in continuous variation (**Fig. 2A**). Even though no distinct segregation between a susceptible and a resistant pool could be seen, a continuous segregation is common and often does result in the identification of QTL when mapped. We therefore continued the genetic analysis.

Bulked segregant analysis (BSA) was performed on a resistant bulk (RS<1.9%; n=18) and a susceptible bulk (**Fig. 2A**; RS>9%; n=18). DNA was isolated, pooled, and bulk sequenced using Illumina short read paired end sequencing. For mapping via k-mer filtering (Prodhomme et al., 2019), the susceptible parent and two resistant genotypes from the R- bulk were also sequenced. A k-mer set enriched in resistance-associated k-mers was obtained and the postitive read pairs from the R-bulk (33724) were mapped to the DM v6.1 genome. In this way we identified 84 significant k-mer bins (exponential distribution, p < 0.05), of which 71 were located on chromosome 11 between 42.37 - 43.72 Mb (**Fig. 2B-C**). The two bins with the highest number of mapped reads were found at 42.83 - 42.85 Mb (**Fig. S3**).

**Fig. 1:**
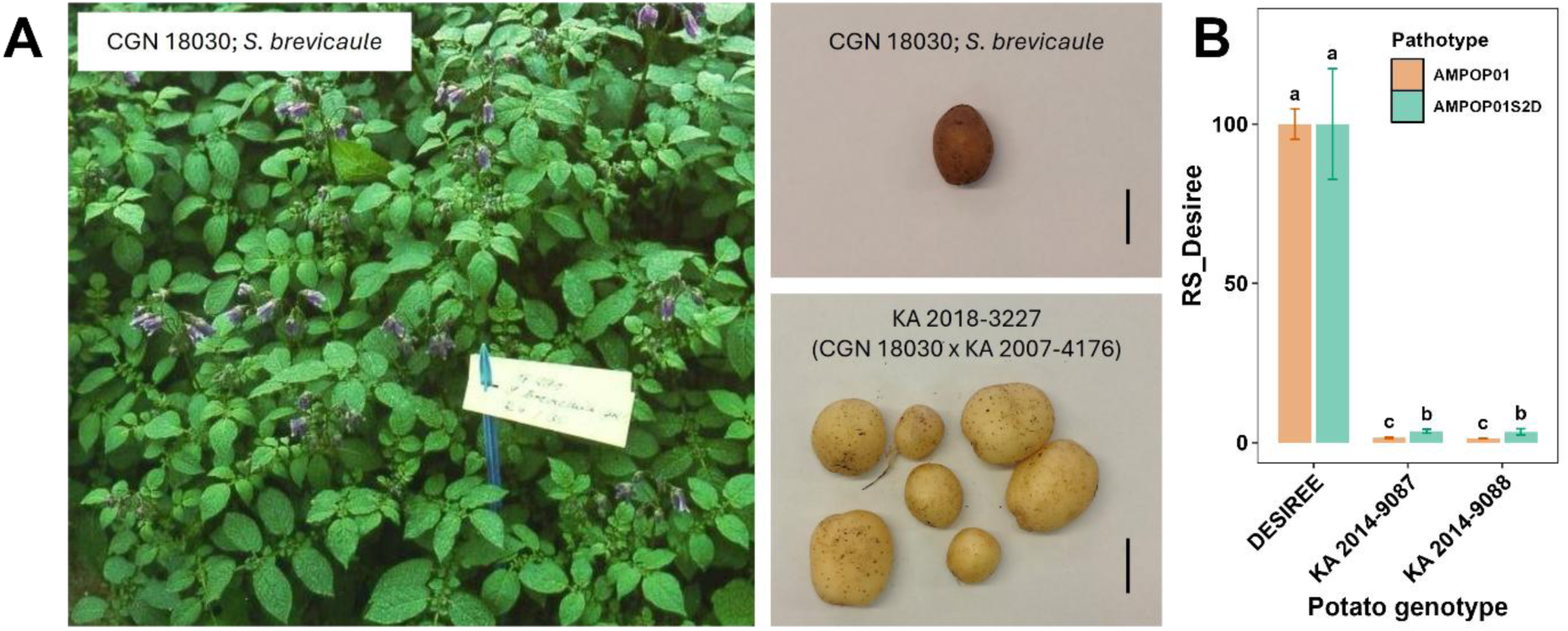
A) Flowering wild potato relative S. brevicaule plants (CGN18030, diploid 2n = 24, 2EBN) originating from Bolivia and tuber yield of one greenhouse pot under long daylight conditions. Bar represents 20 mm (Centre for Genetic Resources (CGN), 1996); Langendoen). B) Two S. brevicaule clones KA 2014-9087 and KA 2014-9088 (CGN 18030) were resistant to different populations of G. pallida. The x-axis shows the different potato genotypes tested in three repetitions and the y-axis shows the relative susceptibility when compared to susceptible genotype Desiree (RS_Desiree). The red bars indicate the results using G. pallida population AMPOP01 (2016), the green bars indicate the results using population AMPOP01S2D (2017). Groups not sharing the same letter differ significantly (p < 0.05, Kruskal-Wallis test, 3 reps. per pathotype/genotype combination).

**Fig. 2:**
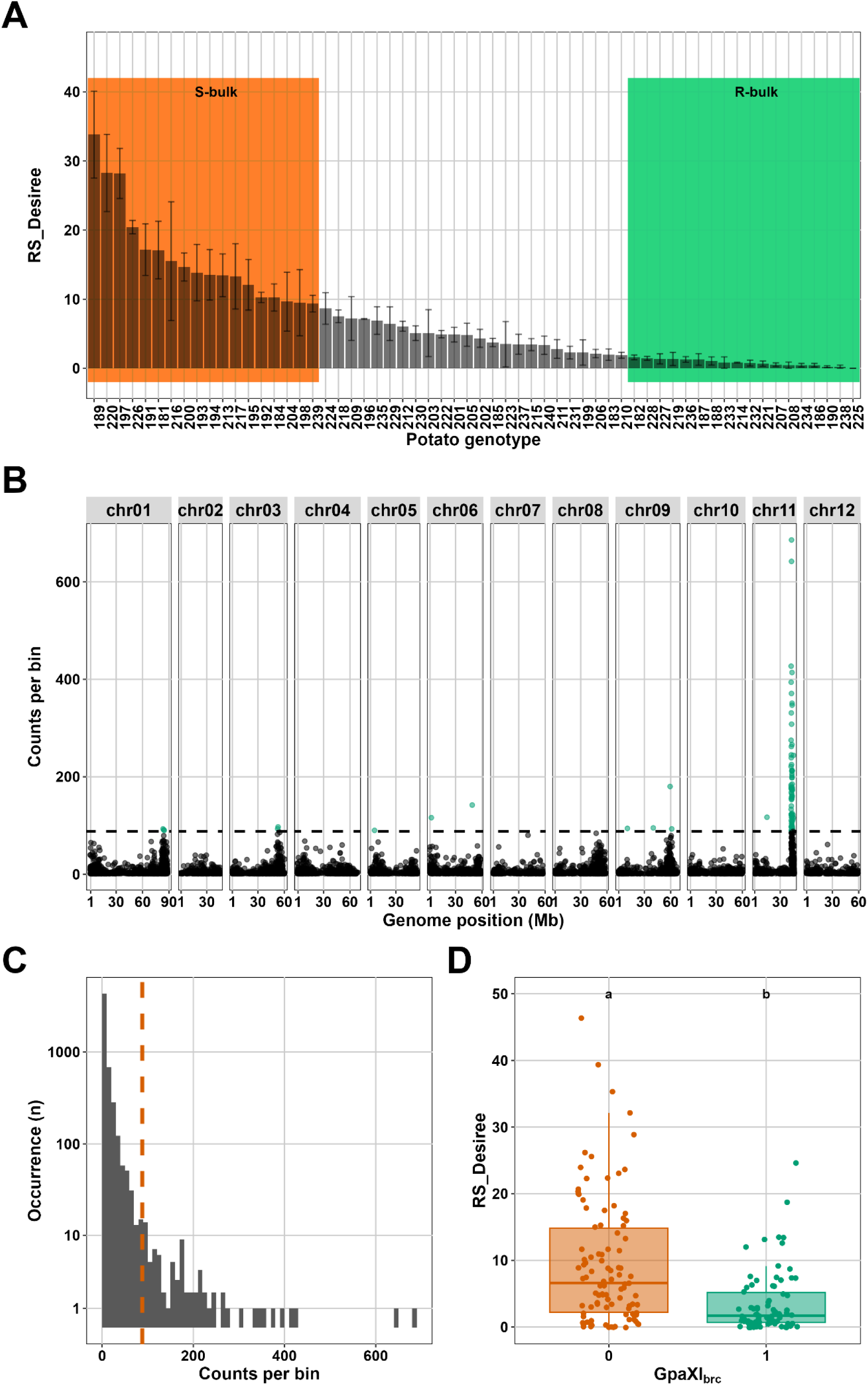
A) Offspring of KA 2014-9087 x susceptible parent (P 2018-3004) segregates for resistance against GpaVvrn-virulent G. pallida. The x-axis shows the potato genotypes from population P2018-3004 and positive control Desiree. The y-axis represents the relative susceptibility when compared to Desiree as tested in a pot test with three replications. Error bars represent the standard error. Genotypes within the orange and green boxes were used to compose the susceptible- and resistant bulk respectively. Genotypes 227 and 228 are KA 2018- 3227 and KA 2018-3228. B) Manhattan plot. The x-axis shows the genomic position on the 12 different chromosomes of S. tuberosum. The y-axis represents the amount of reads per 10kb of reference genome DM v.6.1. Green dots show enriched locations. C) Distribution rate of k-mer counts per bin. Orange line indicates minimal counts to reach significance. D) GpaXIbrc positive potato genotypes score a significantly lower relative susceptibility compared to GpaXIbrc negative potato genotypes in a pot test. The x-axis shows the GpaXIbrc marker score of the different potato genotypes. The y-axis shows the RS compared to Desiree when using G. pallida population AMPOP01. Each genotype was tested in three repetitions, every repetition was plotted (0;N=102, 1;N=78). Groups not sharing the same letter differ significantly (p < 0.05, Kruskal-Wallis test).

We developed a PCR assay to track the resistance marker that is segregating in the mapping population. Primers were selected within and outside the QTL interval on chromosome 11 based on SNPs present in the resistant genotypes and absent in susceptible genotypes. We selected nine primers pairs and tested the effect of the presence or absence of the genetic markers on relative susceptibility of the mapping population, which was analysed using linear regression. SNP location and linear regression results are shown in **Table S5**. Two markers located on the top of chromosome 11 (**Table S4**) did not show a significant correlation with the RS values. Genetic marker Brev4 scored the highest determination coefficient compared to the other markers and was thus selected for marker assisted selection on resistance haplotype *GpaXI_brc_*. The presence of the marker was significantly correlated with resistance (**Fig. 2D**) and segregated roughly in a 1:1 ratio (34:26, χ² = 1.07, df = 1, p ≈ 0.30) as expected.

A list of all annotated genes on potato reference genome DM (v6.1) was retrieved from https://spuddb.uga.edu. It was filtered for location (Chr11.42-46mb) and keyword “resistance”, yielding a list of 46 LRR and/or NB-ARC domain containing disease resistance proteins (**Fig. S5**). Additional fine mapping could be performed to determine whether one of these resistance genes is underlying the *S. brevicaule* resistance, but has limitations and was considered outside of the scope of this study. Instead, we chose to further investigate the agronomic application of the resistance.

### Pyramiding *GpaXIbrc* with other resistance haplotypes increases resistance

To investigate the potential of resistance gene stacking of *GpaXI_brc_* in combination with other *G. pallida* resistance haplotypes, two additional crosses were made. KA 2018-3227 (resistant clone from P 2018-3004) was crossed with *GpaV_vrn_* and *GpaIX_vrn_* donor KA 2015-5546 and *GpaXI_tar_* donor KA 2016-2537 to yield populations P 2020-5501 and P 2018-2006 respectively (**Fig. s1**). After two years of multiplication, tubers of the different genotypes were used in a bioassay. In house genetic markers based on (Tan et al., 2009), (van Eck et al., 2017) and (Bryan et al., 2002; Rouppe van der Voort et al., 2000) were used for identification of *GpaXI_tar_*, *GpaV_vrn_* and *GpaIX_vrn_* respectively. Overall, the genotypes with *GpaXI_brc_* had a significantly lower relative susceptibility compared to the genotypes without *GpaXI_brc_* (PERMANOVA P<0.05) (**Figs. 3 and S6**).

**Fig. 3:**
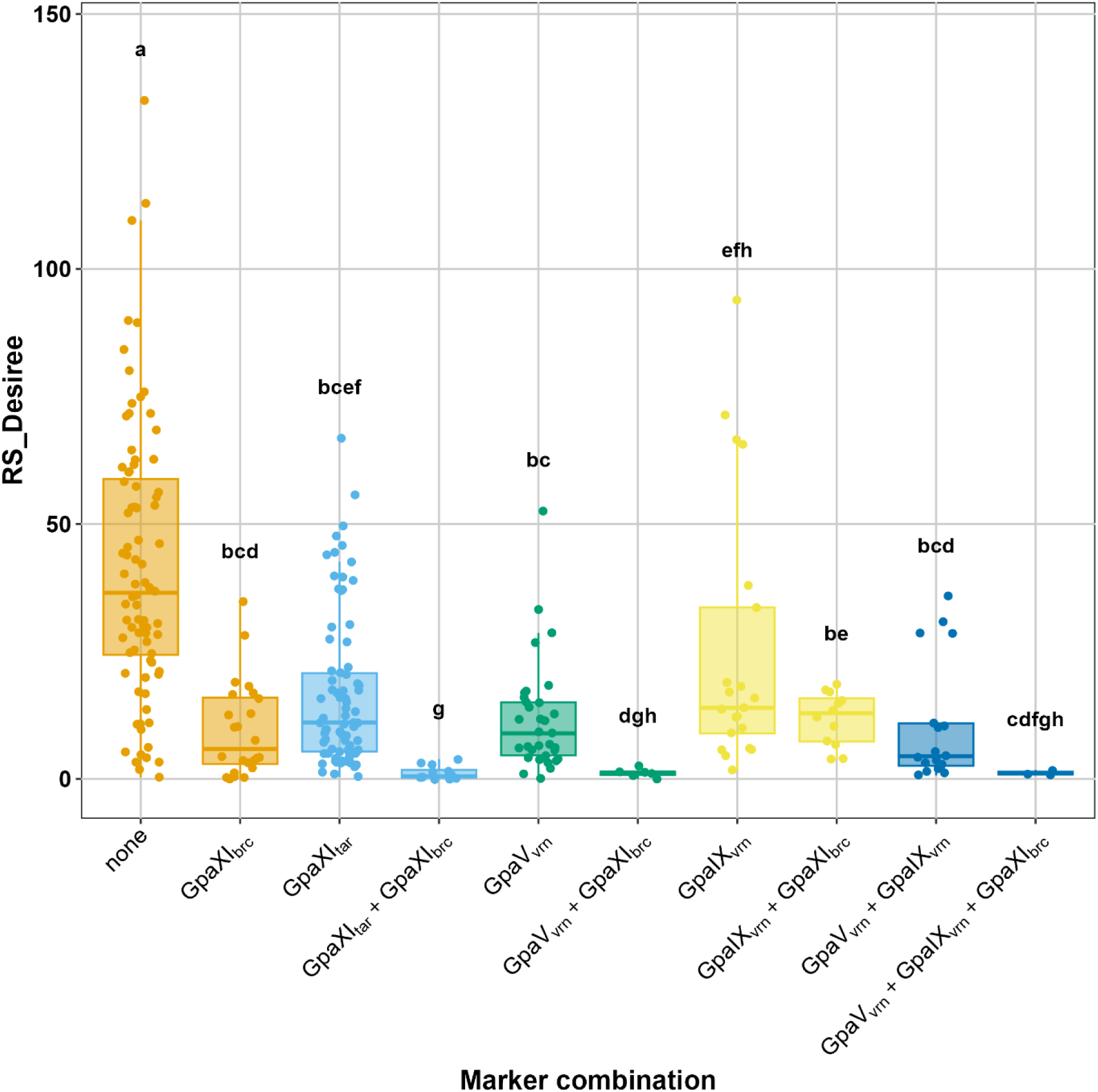
Stacking the new GpaXIbrc marker with GpaVvrn, GpaIXtar or GpaIXvrn further increases resistance against GpaVvrn-virulent G. pallida. The x-axis shows the potato genotypes divided over different marker combinations. The y-axis shows the RS compared to Desiree when using G. pallida population AMPOP01. Each genotype was tested in three repetitions; every repetition was plotted. Groups not sharing the same letter differ significantly (p < 0.05, PERMANOVA).

Although in general the number of individuals containing stacks of *GpaXI_tar_*, *GpaV_vrn_* or *GpaIX_vrn_* with *GpaXI_brc_* was low, significant differences were found between the control group without resistance genes (N=28 genotypes) and the group with *GpaXI_brc_* (N=8 genotypes), the *GpaXI_tar_* group (N=27 genotypes) and the *GpaXI_tar_* + *GpaXI_brc_* group (N=4 genotypes) and the *GpaVvrn* group (N=11 genotypes) and the *GpaVvrn* + *GpaXIbrc* group (N=2 genotypes). No significant differences were found between groups *GpaIXvrn* (N=7 genotypes) and *GpaIXvrn* + *GpaXIbrc* (N=4 genotypes) and between *GpaVvrn* + *GpaIXvrn* (N=6 genotypes) and *GpaVvrn* + *GpaIXvrn* + *GpaXIbrc* (N=1 genotype). The latter could potentially be explained by the low number of individuals within the *GpaVvrn* + *GpaIXvrn* + *GpaXIbrc* group. Stacking of *GpaXIbrc* to other resistance genes thus increases resistance against *GpaVvrn*-virulent *G. pallida*.

A PERMANOVA was used to test whether any of the combinations lead to a non-additive effect, indicating that the combination of two markers yielded a higher resistance than the sum of two individual markers. The combination of *GpaXI_brc_* with *GpaXI_tar_* was significantly more resistant to PCN than the separate markers (p = 0.03), but the effect was minor (R^2^ = 0.015). The other combinations were found to be additive (**Table S6-7**). However, a trend is visible that stacks of *GpaXI_brc_* with other resistance haplotypes (apart from *GpaIX_vrn_*) perform better than expected based on their individual resistance levels (positive epistasis).

### Characterization of the *GpaXIbrc* resistance mechanism *in vitro*

To characterize the resistance mechanism underlying the *GpaXI_brc_* haplotype, an *in vitro* bioassay was performed. KA 2020-2084, containing the *GpaXI_brc_* marker, was compared to susceptible *S. brevicaule* clone P 2020-5501-44 and two potato cultivars Desiree (susceptible) and Aletta (resistant (Tammes JE & Langendoen D, 2025)).

At 5 dpi, all potato genotypes showed browning and necrosis of root tissue, which is a sign of migration damage. PCN could thus infiltrate root tissue of all genotypes. Aletta showed notably less migration damage than the other genotypes.

At 28 dpi, adult males were visible either still encapsulated attached to the root or as an adult on the agar in search for a female. Also, adult females could be counted (**Table 1**). Desiree had significantly more adult males and adult females when compared to the other genotypes.

**Table 1:**
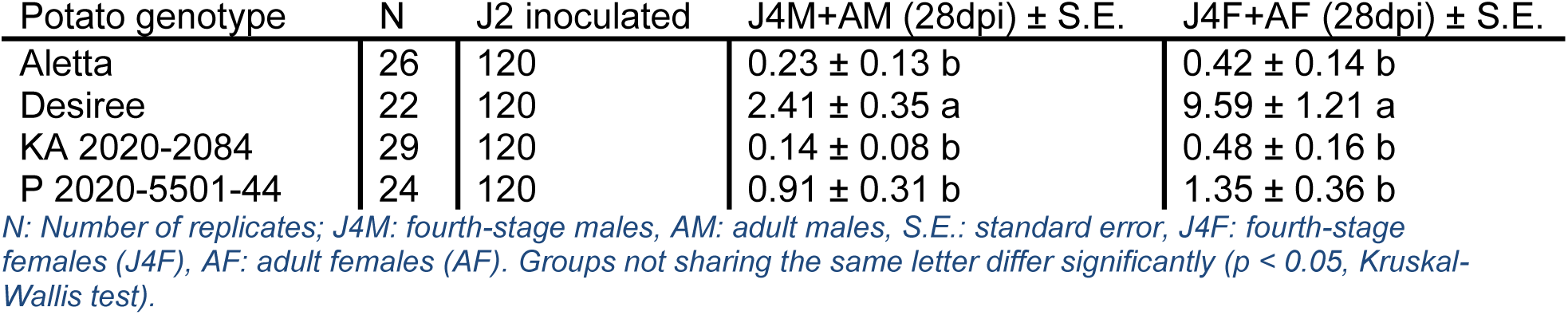
Susceptibility of potato genotypes to in vitro Globodera pallida infection at 28 days post-inoculation (dpi).

At 63 dpi, on Desiree roots, many fully developed cysts were found, often on thickened root fragments, indicating proper feeding site formation (**Fig. 4**). In Aletta, few cysts were found, but no discoloration of the roots or any other symptoms of infection or invasion. P 2020- 5501-44 unexpectedly showed few cysts in contrast to the *in vivo* bioassays. Many roots were completely discolored, which could indicate dead roots. KA 2020-2084, which contains the *GpaXI_brc_* marker, showed few cysts. In some roots small necrotic patches could be observed, potentially indicating programmed cell death by a hypersensitive response. No male PCN could be observed in any genotypes.

**Fig. 4:**
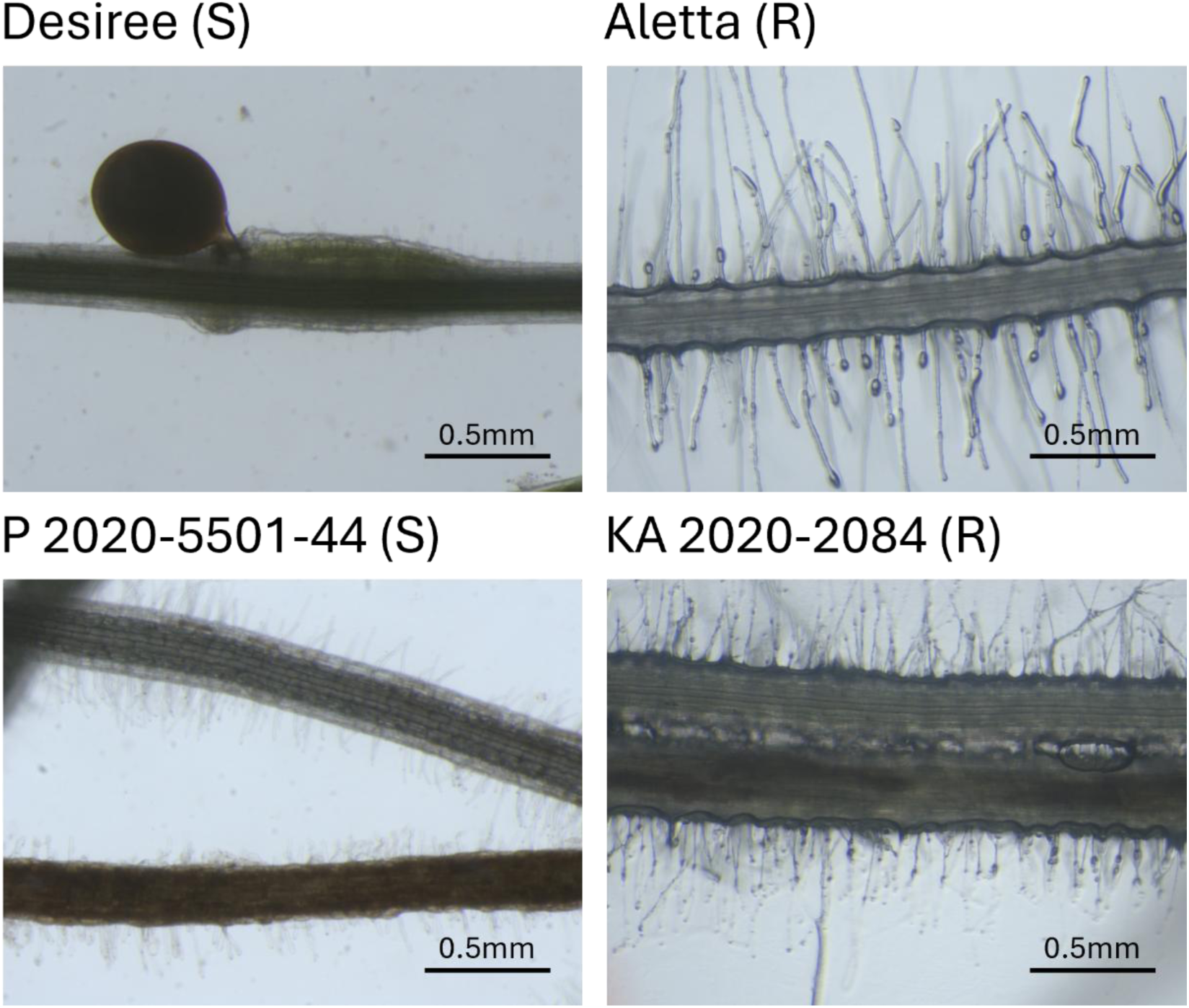
In vitro potato roots inoculated with Globodera pallida AMPOP01. Each panel shows representative image of the roots of each potato genotype. (S)=susceptible, (R)=resistant. Scale bars represent 0.5 mm.

## Discussion

### A *S. brevicaule* resistance mapping to Chromosome 11

After screening for resistance against *GpaV_vrn_*-virulent *G. pallida*, a novel source of resistance was found in wild potato relative *S. brevicaule*. It was crossed with a susceptible diploid *S. tuberosum* to create a mapping population that could be used to identify the location of the resistance causing haplotype. The mapping population showed continuous variation when phenotyped for *G. pallida* resistance and the most- and least resistant individuals were bulked before Illumina sequencing alongside the associated susceptible and resistant genotypes. Resistance-linked 31 bp k-mers were mapped to the *S. tuberosum* reference genome, yielding a large peak on the bottom of chromosome 11. Because it was difficult to design unique primers within the highly homogeneous resistance gene clusters (Jacobs et al., 2010), primers were developed around a resistance linked SNP (Chr11. 43Mb) between these clusters. Additional finemapping would be required to determine the exact resistance gene(s) underlying the *S. brevicaule* resistance. The marker segregated in a 1:1 ratio and was significantly correlated with resistance. The newly developed genetic marker was named *GpaXI_brc_*.

The *GpaXI_brc_*-resistance maps to an NB-LRR rich locus. Other *Globodera* resistance loci map to the approximate location of *GpaXI_brc_* (Gartner et al., 2021), such as Gro1.3 (Kreike et al., 1993) and *GpaXI_spl_* (Caromel et al., 2005). *GpaXI_tar_* does map to Chr11, but it is located on the other arm as *GpaXI_brc_*. In *S. sparsipilum*, the stacking of minor resistance gene *GpaXI_spl_* to major resistance gene *GpaV_spl_* changes the resistance phenotype from masculinisation of the penetrated larva to rapid necrotic reaction (Fournet, 2022; Kerlan et al., 2017). We found no masculinisation response in our *GpaXI_brc_* harbouring genotype but did find localized necrotic regions that could indicate a hypersensitive response. Therefore, potentially, it is a different resistance haplotype than *GpaXI_spl_*. However, because neither gene has been cloned, we cannot rule out these two haplotypes are in fact the same resistance gene. For this purpose, the newly published *S. brevicaule* genome (Shanmugavel et al., 2026) could be of help, since the *S. tuberosum* reference genome DM6.1 probably does not contain the *GpaXI_brc_* haplotype.

The *GpaXI_brc_* resistance is a small-effect resistance. Although *GpaXI_brc_* was the marker that explained most of the variance in the mapping population, it only accounted for 14.6 % of the variance. Therefore, we speculate that other loci conferring resistance may be found in *S. brevicaule* that underly the considerable remainder of the variance in resistance. Many *Globodera* resistance QTL mapped to date have a small effect on the resistance level of the plant. For instance, resistance against *G. rostochiensis* originating from *S. spegazzinii* (Kreike et al., 1993), *G. pallida* resistance originating from *S. sparsipilum* (Caromel et al., 2005), and QTL like *GpaIX_vrn_* from *S. vernei*. In retrospect, additional small-effect resistance genes in *S. brevicaule* may potentially be found in up to four additional chromosome regions, with less strong BSA signals, that were visible in the intermediate steps of the k-mer filtering process. However, because these regions could also be false positives, these were not explored.

From a resistance breeding point of view, the need for many different small-effect resistance genes to provide total protection of the plant makes breeding challenging. However, it also drives diversification of the resistances in cultivars, which may in-turn improve durability and thereby the agronomic lifespan of the resistance.

### Pyramiding *GpaXIbrc* with other resistance haplotypes increases resistance

Stacking resistance genes generally leads to a higher resistance level against PCN (Dalton et al., 2013; Price et al., 2024; Rigney et al., 2017). We stacked the novel *S. brevicaule* resistance with known resistance haplotypes *GpaXI_tar_*, *GpaV_vrn_*, and *GpaIX_vrn_*. The mean relative susceptibility of genotypes with *GpaXI_brc_* was significantly lower than the mean relative susceptibility of genotypes without *GpaXI_brc_*. Stacking of *GpaXI_brc_* to *GpaXI_tar_* and *GpaV_vrn_* significantly lowered the relative susceptibility. A similar trend was visible when *GpaXI_brc_* was combined with *GpaIX_vrn_* and a combination of *GpaIX_vrn_* and *GpaV_vrn_*, yet the limited representation of genotypes exhibiting these specific combinations probably hampered the identification of statistically significant differences.

Stacks of resistance haplotypes in this study (with the exception of *GpaXI_brc_* + *GpaIX_vrn_*) performed better than expected based on the resistance effect of separate resistance QTL. However, only in combination *GpaXI_tar_* + *GpaXI_brc_*, this effect was significant, potentially because of the low number of unique recombinants, and therefore replicates, obtained per combination. An explanation for this positive epistasis could be that the two resistance haplotypes target different steps in the infection process of *G. pallida*. Overcoming both resistance haplotypes simultaneously requires that *G. pallida* possesses multiple virulence alleles, which is less likely than possessing a single virulence allele overcoming a single haplotype (e.g. as in *Gp-pat-1*).

Although studies that follow stacks of resistance genes over multiple years in potato are rare, some show that stacks of resistance genes/haplotypes lead to durable resistance. For instance, a stack of three late blight resistance genes in potato provided complete resistance in the field over several seasons (Ghislain et al., 2019). In other crops, the stacking of minor resistance genes with major resistance genes was found to increase their durability as reviewed by (Mundt, 2018). Models, on the other hand, show that durability is increased upon stacking, but selection on multiple resistance genes could lead to multi virulent populations (Crété et al., 2020). Therefore, it is crucial to keep monitoring and evaluating the infestation in the field and to include other disease management practices, such as crop rotation, soil management and targeted (biological) control (Sikora et al., 2021).

### Breeding with *S. brevicaule*

Although many resistance genes have been described, resistance in modern potato cultivars has a small genetic base (mainly *GpaV_vrn_* in Europe (Schaveling, van Rijt, et al., 2026)). The molecular markers published in this study are valuable for potato breeders to select genotypes with novel *GpaXI_brc_* resistance, as it is almost impossible to breed for stacks of quantitative resistance traits, based solely on phenotyping (Leuenberger et al., 2025). The fact that no major resistance genes map to the same location as *GpaXI_brc_* (Gartner et al., 2021) drastically improves its use in breeding. Potato cultivars with an RS for *G. pallida* of <1% are indicated as highly resistant in the Dutch variety catalogue. Some genotypes with stacks of resistance haplotypes including *GpaXI_brc_* reach this threshold, making *GpaXI_brc_* extra valuable.

Breeding with *S. brevicaule* material can have added benefits, besides its *G. pallida* resistance. The progenitor *S. brevicaule,* CGN18030, is also resistant or less susceptible to warts disease, gangrene and potato virus X (PVX) (Centre for Genetic Resources (CGN), 1996). Although the CGN reports it to be susceptible to late blight, late blight resistance is highly dependent on the *Phytophthora infestans* strain in the field (Deng et al., 2025). Since other researchers (Enciso-Maldonado et al., 2022) reported another *S. brevicaule* (PI 631209; VIR 7370) to be resistant to *P. infestans*, it could be worthwhile to test the material used in this study for late blight resistance. Lastly, *S. brevicaule* has also been reported to be resistant to soft rot (Chung et al., 2010) highlighting its potential.

## Conclusion

In this study we map a resistance locus in wild potato relative *Solanum brevicaule* to the bottom of chromosome 11 (42.83 - 42.85 Mb) using a k-mer based bulked segregant analysis and name it *GpaXI_brc_.* The presence of the *GpaXI_brc_* marker in the potato genome significantly increases resistance against a GpaV_vrn_-virulent *G. pallida* population (AMPOP01). Furthermore, the *GpaXI_brc_* marker was stacked with three other resistance haplotypes: *GpaXI_tar_*, *GpaV_vrn_* and *GpaIX_vrn_*. Stacking of the novel *GpaXI_brc_* haplotype resulted in a significant increase in resistance. The largest effect was seen in combinations with “major” resistance haplotypes *GpaV_vrn_* + *GpaIX_vrn_* and *GpaXI_tar_* + *GpaXI_brc_*. To determine we are dealing with a novel gene, the resistance mechanism underlying the *GpaXI_brc_* haplotype was described after performing an *in vitro* bioassay. The resistance mechanism was different from *GpaXI_spl_* as described in literature, which is also localized at the bottom of chromosome 11. Therefore, we conclude that we found a novel *G. pallida* resistance haplotype originating from *S. brevicaule*.

## Statements & Declarations

### Author contributions

Design of breeding schemes was done by ANOR and JET. Greenhouse bioassays for resistance testing were performed by DL, HR and AW. The bulked segregant analysis was done by JMdB, RHZ and MGS. HR and IM developed genetic markers. The *in-vitro* bioassays were conducted by YD, J-JW and DL. Manuscript was written by DL under supervision of MGS. All authors provided feedback on the draft.

## Acknowledgements

The authors are grateful to Geert Smant, Aska Goverse and Johan Hopman for their guidance in supervision and discussion. We also extend sincere thanks to Marten Kemper, Diana Mulder-Grooten and Liesbeth Stolker for performing the crosses required for this study. Further thanks go to Mick Wardenburg, Natasja Dammer and Harm van Emst for their work in testing genetic markers and to Paul Heeres and Els Heeres for early PCN tests.

Finally, we thank Jan van der Heide, Allard Bakema, Ties Drent, Frank van Boven, Sander Elting and Titus Kamping for propagating the potatoes in both field and greenhouse settings.

---

**Fig. S1:**
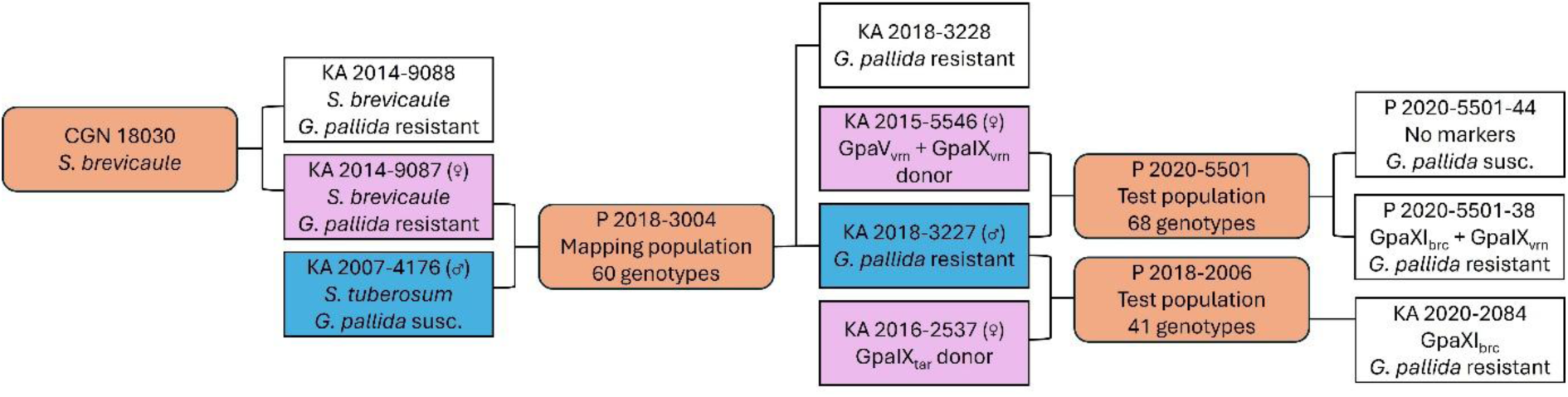
Population pedigree of material used in this research. Rectangles (white (not crossed), pink (crossed as female) or blue (crossed as male)) represent individual genotypes; ellipses (orange) represent grouped individuals.

**Fig. S2:**
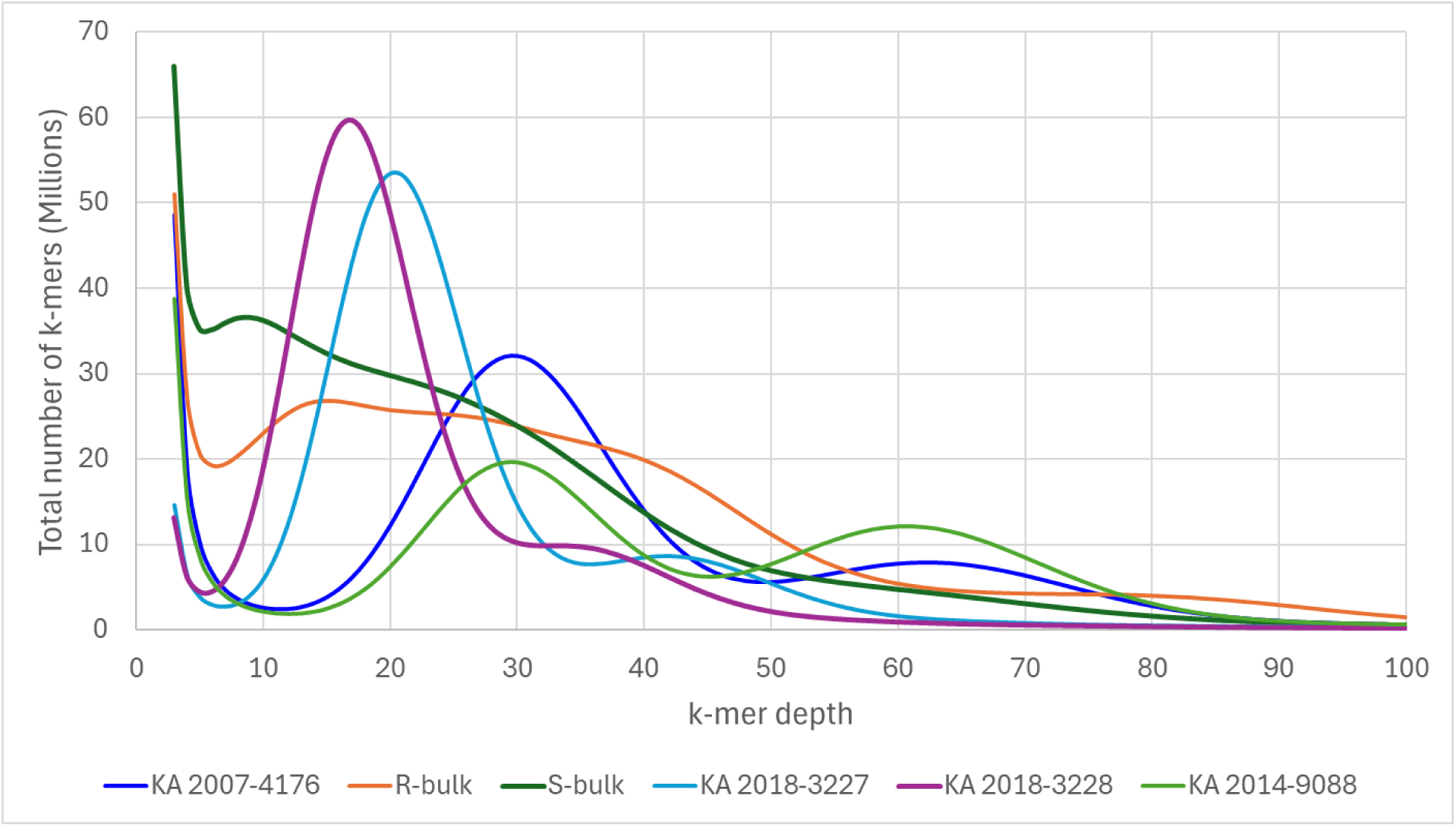
Distribution of k-mers in different samples.

**Fig. S3:**
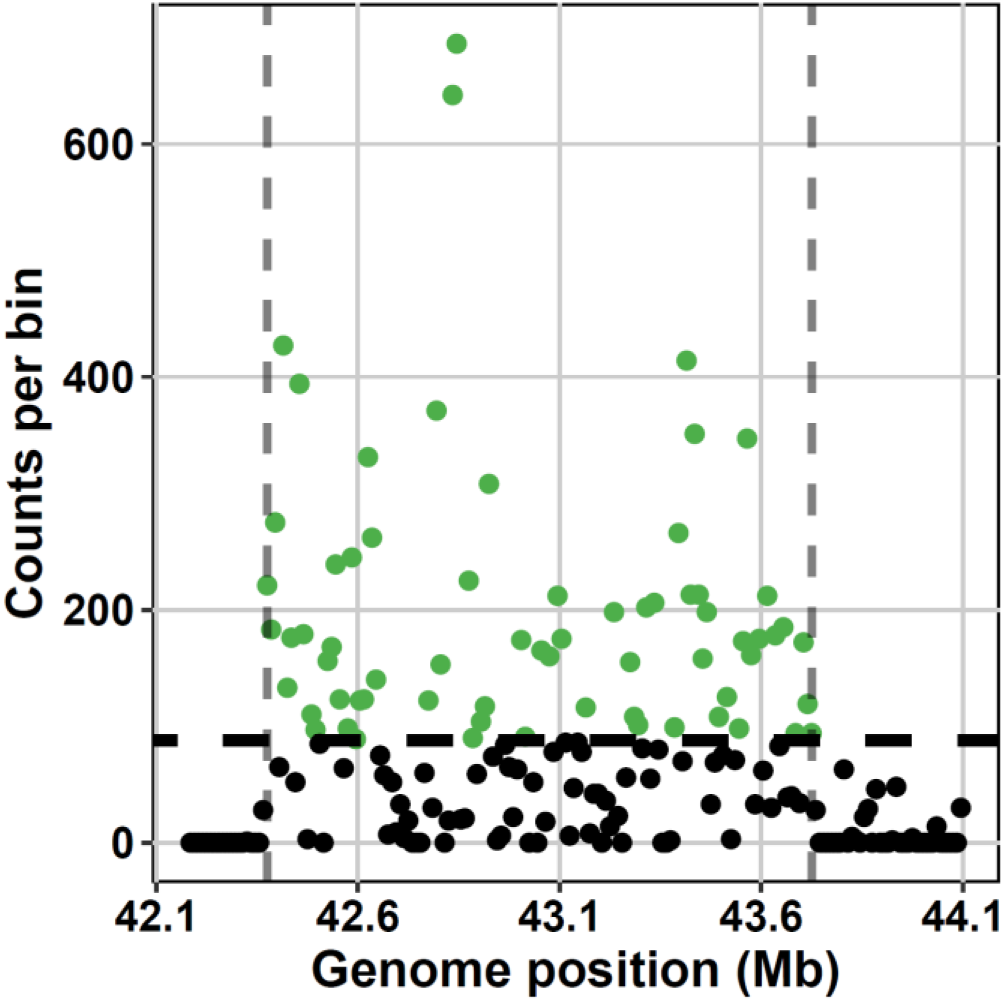
GpaXIbrc locus on chromosome 11. The x-axis shows the genomic position chromosome 11 of S. tuberosum. The y-axis represents the amount of reads per 10kb of reference genome DM v.6.1. Green dots show enriched locations.

**Fig. S4:**
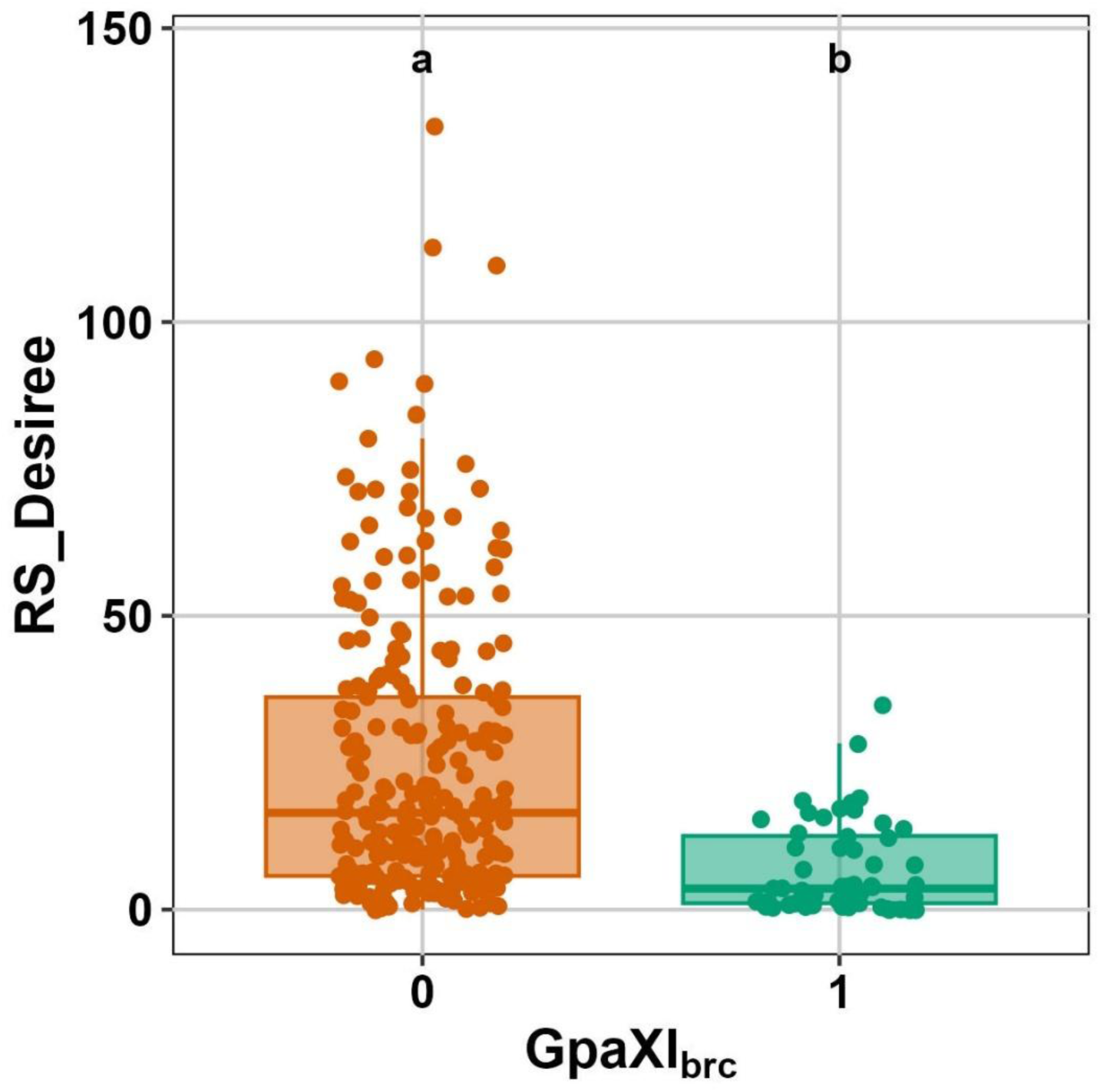
Effect of GpaXIbrc in pyramiding experiment. Presence of GpaXIbrc increases resistance to G. pallida AMPOP01 in addition to the effect of other resistance loci. Groups not sharing the same letter differ significantly (p < 0.05, Kruskal-Wallis test).

**Fig. S5:**
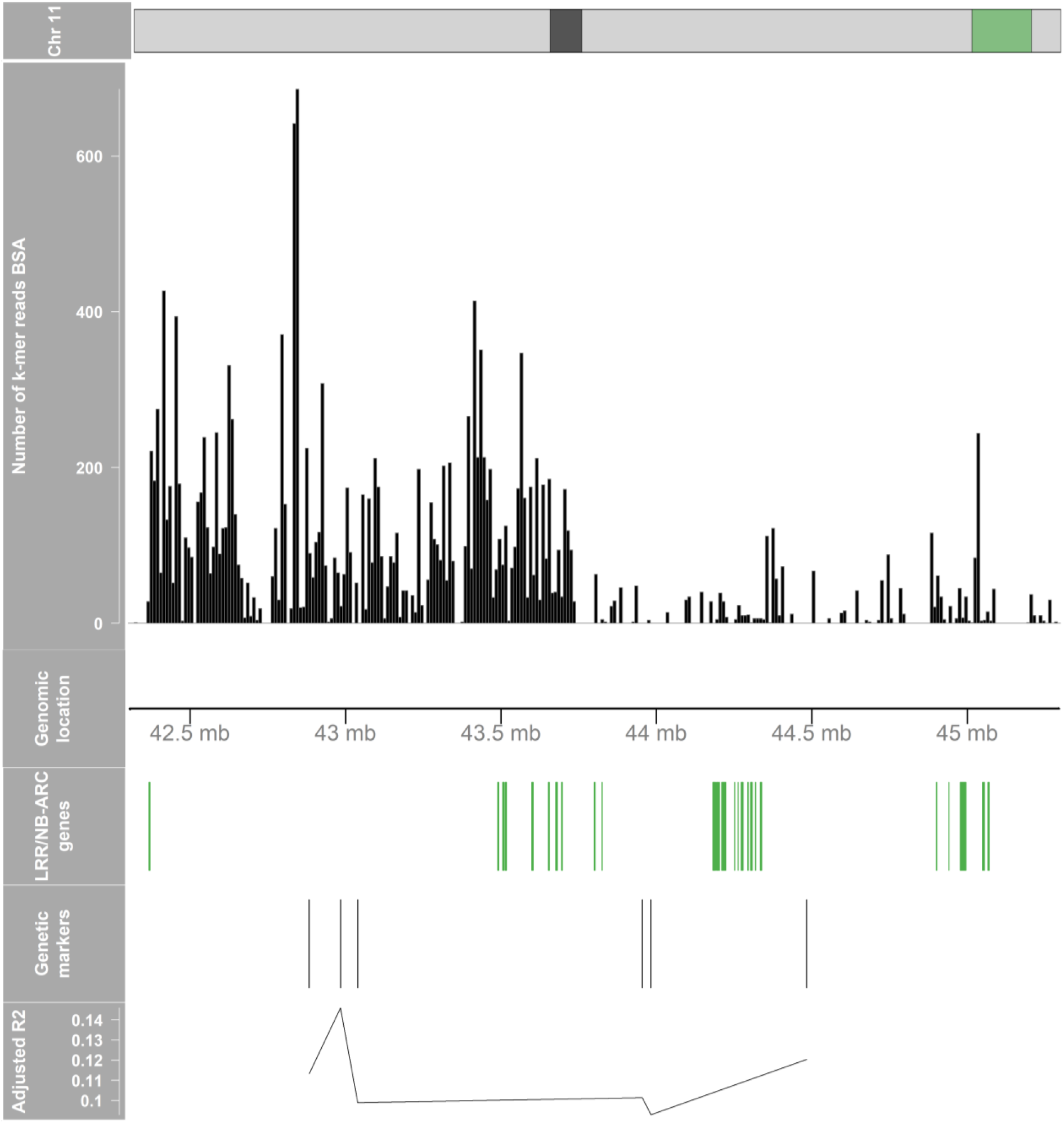
Combined plot of BSA results, genomic location, LRR/NB-ARC genes, genetic markers and their R^2^adj. Top track indicates the Chr11.42-46mb position in green, with the centromere in black. Second track shows the BSA results. Third track shows the genomic location of chromosome 11. Fourth track shows the LRR/NB-ARC genes as retrieved from https://spuddb.uga.edu/. It was filtered for location (Chr11.42-46mb) and keyword “resistance”, yielding a list of 46 LRR and/or NB-ARC domain containing disease resistance proteins. Fifth track shows the location of the Chr11.42-46mb markers used in this study. The last track shows the adjusted R^2^ of the genetic markers as obtained by linear regression.

**Table S1:** Resistance QTL in material used in this research.

| Genotype | ♀ | ♂ | Grp1<sup>a</sup> | GpaXI<sub>tar</sub><sup>b</sup> | GpaIV<sub>and</sub><sup>c</sup> | GpaV<sub>vrn</sub><sup>d</sup> | GpaIX<sub>vrn</sub><sup>e</sup> | GpaV<sub>spl</sub><sup>f</sup> | GpaXI<sub>brc</sub><sup>g</sup> |
|---|---|---|---|---|---|---|---|---|---|
| KA 2014-9088 | CGN18030 (brc) | CGN18030 (brc) | not tested | not tested | 0 | 0 | 0 | 0 | 1 |
| KA 2014-9087 | CGN18030 (brc) | CGN18030 (brc) | not tested | not tested | 0 | 0 | 0 | not tested | 1 |
| KA 2007-4176 | IVP 04-97-2 | STET 96-2039-10 | 0 | 0 | 0 | 0 | 0 | 0 | 0 |
| KA 2018-3228 | KA 2014-9087 | KA 2007-4176 | 0 | 0 | 0 | 0 | 0 | 0 | 1 |
| KA 2015-5546 | KA 2009-5916 | KA 2007-4174 | 0 | 0 | 0 | 1 | 1 | 0 | 0 |
| KA 2018-3227 | KA 2014-9087 | KA 2007-4176 | 0 | 0 | 0 | 0 | 0 | 0 | 1 |
| KA 2016-2537 | KA 2010-2261 | KA 2010-4288 | 0 | 1 | 0 | 0 | 0 | not tested | 0 |
| P 2020-5501-44 | KA 2015-5546 | KA 2018-3227 | 0 | 0 | 0 | 0 | 0 | 0 | 0 |
| P 2020-5501-38 | KA 2015-5546 | KA 2018-3227 | 0 | 0 | 0 | 0 | 1 | 0 | 1 |
| KA 2020-2084 | KA 2016-2537 | KA 2018-3227 | 0 | 0 | 0 | 0 | 0 | 0 | 1 |

*a) (Finkers-Tomczak et al., 2009; Rouppe van der Voort et al., 2000)*
*b) (Tan et al., 2009)*
*c) (Bradshaw et al., 1998)*
*d) (Rouppe van der Voort et al., 2000; van Eck et al., 2017)*
*e) (Bryan et al., 2002; Rouppe van der Voort et al., 2000)*
*f) (Caromel et al., 2005)*
*g) this study*

**Table S2:** Sequence statistics for the BSA WGS samples.

| Genotype / sample | Description | Ploidy | Estimated sequence depth (a) |  |
|---|---|---|---|---|
|  |  |  | Sequence reads | k-mer database (b) |
| KA 2014-9087 | Resistant mother (c) | 2x | n/a | n/a |
| KA 2007-4176 | Susceptible father | 2x | 99 | 58 |
| R-bulk | Pool of 18 resistant F1 individuals | n/a | 115 | 67 |
| S-bulk | Pool of 18 susceptible F1 individuals | n/a | 97 | 57 |
| KA 2018-3227 | Individual from R-BULK | 2x | 62 | 36 |
| KA 2018-3228 | Individual from R-BULK | 2x | 53 | 31 |

*(a) based on number of 150 bp reads and a genome size of 750 Mbp*
*(b) calculated for 31 bp k-mers extracted from 150 bp reads*
*(c) genotype was no longer available at the time of sequencing*

**Table S3:** steps in the BSA k-mer filtering procedure.

| BSA step | K-mer database stage | Number of k-mers (a) |
|---|---|---|
| 0 | R-bulk | 1563137415 |
| 1 | subtract S-bulk | 230547936 |
| 2 | subtract KA 2007-4176 | 208198601 |
| 3 | intersect KA 2018-3227 (b) | 5615012 |
| 4 | intersect KA 2018-3228 | 1768440 |
| 5 | reduce to simplex k-mers (depth 10-30) in KA 2018-3227 | 751878 |

*(a) k-mer depth values of 2 or more*
*(b) the k-mer depth values were changed to those for KA 2018-3227*

**Table S4:**
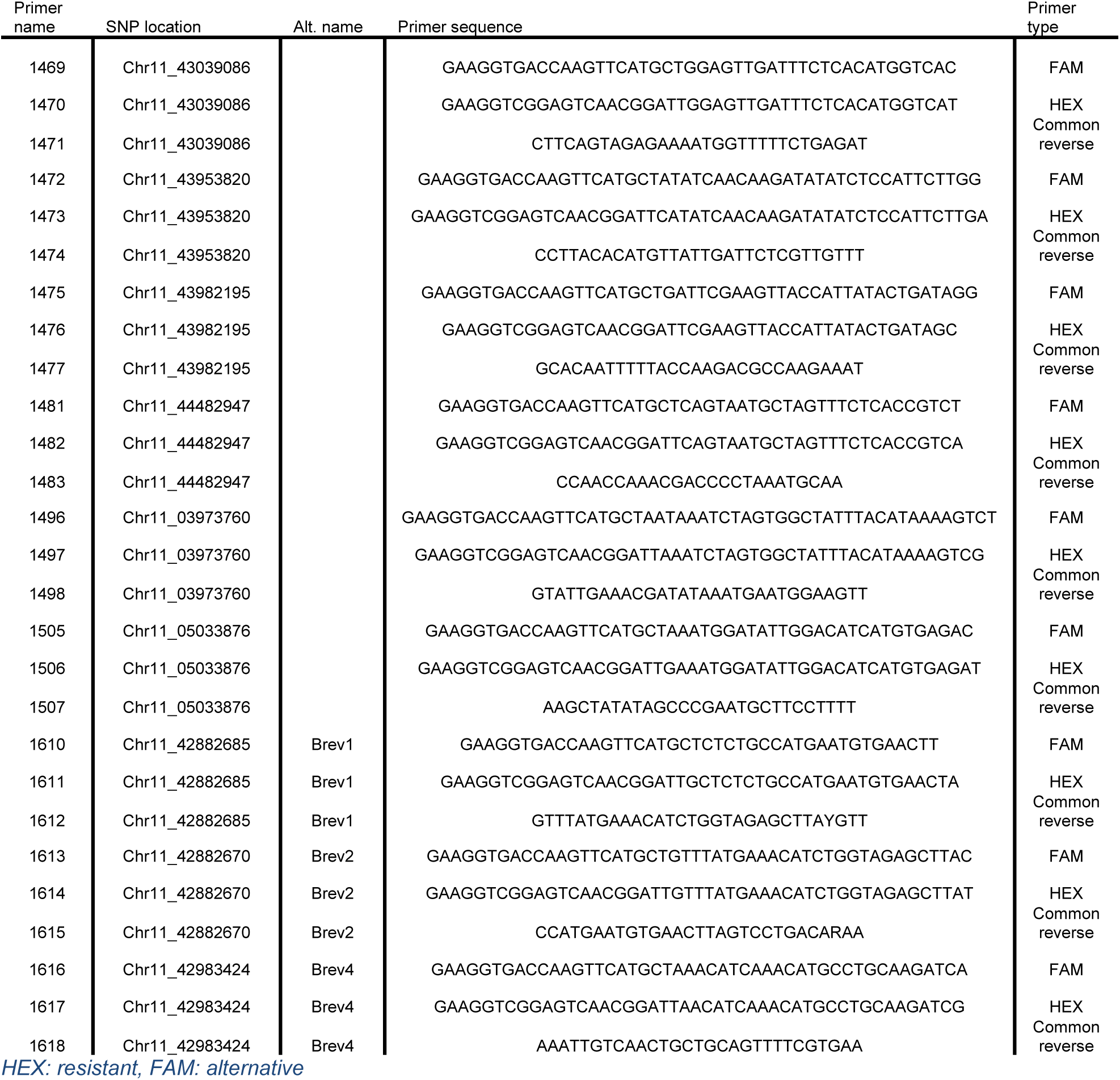
Primer sequences for PACE genetic marker assays.

**Table S5:** Genetic marker location and linear regression results.

| Name | Location (bp) | P-value | Adjusted R<sup>2</sup> |
|---|---|---|---|
| 1496*1497*1498 | Chr11.3973760 | 0.2855 | 0.004574 |
| 1505*1506*1507 | Chr11.5033876 | 0.9391 | -0.01912 |
| Brev2 | Chr11.42882670 | 0.006037 | 0.1133 |
| Brev1 | Chr11.42882685 | 0.01114 | 0.09369 |
| Brev4 (*GpaXI<sub>brc</sub>*) | Chr11.42983424 | 0.002337 | 0.1459 |
| 1469*1470*1471 | Chr11.43039086 | 0.00877 | 0.09893 |
| 1472*1473*1474 | Chr11.43953820 | 0.009075 | 0.1014 |
| 1475*1476*1477 | Chr11.43982195 | 0.01349 | 0.09295 |
| 1481*1482*1483 | Chr11.44482947 | 0.004738 | 0.1204 |

**Table S6:** (non-)additive effect of marker combinations on resistance against G. pallida as analysed by PERMANOVA.

|  | R<sup>2</sup> |  |  |  |  |  |  | p |  |  |  |  |  |  |
|---|---|---|---|---|---|---|---|---|---|---|---|---|---|---|
| **Combination (A+B+C)** | **A** | **B** | **A:B** | **C** | **A:C** | **B:C** | **A:B:C** | **A** | **B** | **A:B** | **C** | **A:C** | **B:C** | **A:B:C** |
| *GpaXI<sub>tar</sub>* + *GpaXI<sub>brc</sub>* | 0,164 | 0,164 | 0,015 |  |  |  |  | 0,001 | 0,001 | 0,030 |  |  |  |  |
| *GpaV<sub>vrn</sub>* + *GpaXI<sub>brc</sub>* | 0,166 | 0,176 | 0,018 |  |  |  |  | 0,001 | 0,001 | 0,059 |  |  |  |  |
| *GpaIX<sub>vrn</sub>* + *GpaXI<sub>brc</sub>* | 0,050 | 0,185 | 0,018 |  |  |  |  | 0,005 | 0,001 | 0,065 |  |  |  |  |
| *GpaV<sub>vrn</sub>* + *GpaIX<sub>vrn</sub>* | 0,231 | 0,027 | 0,014 |  |  |  |  | 0,001 | 0,026 | 0,098 |  |  |  |  |
| *GpaV<sub>vrn</sub>* + *GpaIX<sub>vrn</sub>* + *GpaXI<sub>brc</sub>* | 0,151 | 0,028 | 0,006 | 0,146 | 0,013 | 0,012 | 0,003 | 0,001 | 0,005 | 0,184 | 0,001 | 0,054 | 0,064 | 0,346 |

**Table S7:** (non-)additive effect vs. expected.

| Group | Mean RS | Fraction of "none" | Exp. additive RS |
|---|---|---|---|
| none | 41,8 | 1,00 |  |
| *GpaXI<sub>brc</sub>* | 9,6 | 0,23 |  |
| *GpaXI<sub>tar</sub>* | 16,6 | 0,40 |  |
| *GpaXI<sub>tar</sub>* + *GpaXI<sub>brc</sub>* | 1,2 | 0,03 | 3,81 |
| *GpaV<sub>vrn</sub>* | 11,8 | 0,28 |  |
| *GpaV<sub>vrn</sub>* + *GpaXI<sub>brc</sub>* | 1,2 | 0,03 | 2,71 |
| *GpaIX<sub>vrn</sub>* | 25,4 | 0,61 |  |
| *GpaIX<sub>vrn</sub>* + *GpaXI<sub>brc</sub>* | 11,8 | 0,28 | 5,82 |
| *GpaV<sub>vrn</sub>* + *GpaIX<sub>vrn</sub>* | 10,4 | 0,25 |  |
| *GpaV<sub>vrn</sub>* + *GpaIX<sub>vrn</sub>* + *GpaXI<sub>brc</sub>* | 1,2 | 0,03 | 2,38 |

## References

1. Beniers, J. E., Nöllen, Y., van Eck, H. J., & Schouten, H. J. (2019). Increased virulence of Globodera pallida during repeated rearing on different resistant potato cultivars explained by a simple model. Plant Pathology, 68(3), 445–453. 10.1111/ppa.12969

2. Bradshaw, J. E., Hackett, C. A., Meyer, R. C., Milbourne, D., McNicol, J. W., Phillips, M. S., & Waugh, R. (1998). Identification of AFLP and SSR markers associated with quantitative resistance to Globodera pallida (Stone) in tetraploid potato (Solanum tuberosum subsp. tuberosum) with a view to marker-assisted selection. Theoretical and Applied Genetics, 97(1), 202–210. 10.1007/s001220050886

3. Bryan, G., McLean, K., Bradshaw, J., De Jong, W., Phillips, M., Castelli, L., & Waugh, R. (2002). Mapping QTLs for resistance to the cyst nematode Globodera pallida derived from the wild potato species Solanum vernei. Theoretical and Applied Genetics, 105(1), 68–77. 10.1007/s00122-002-0873-9

4. Caromel, B., Mugniéry, D., Kerlan, M.-C., Andrzejewski, S., Palloix, A., Ellissèche, D., Rousselle-Bourgeois, F., & Lefebvre, V. (2005). Resistance Quantitative Trait Loci Originating from Solanum sparsipilum Act Independently on the Sex Ratio of Globodera pallida and Together for Developing a Necrotic Reaction. Molecular Plant-Microbe Interactions MPMI, 18(11), 1186–1194. 10.1094/MPMI

5. Centre for Genetic Resources (CGN), the N. (1996, February 27). Solanum brevicaule. 10.18730/1KJEF

6. Chung, Y. S., Holmquist, K., Spooner, D. M., & Jansky, S. H. (2010). A Test of Taxonomic and Biogeographic Predictivity: Resistance to Soft Rot in Wild Relatives of Cultivated Potato. Phytopathology®, 101(2), 205–212. 10.1094/PHYTO-05-10-0139

7. Crété, R., Pires, R. N., Barbetti, M. J., & Renton, M. (2020). Rotating and stacking genes can improve crop resistance durability while potentially selecting highly virulent pathogen strains. Scientific Reports, 10(1). 10.1038/s41598-020-76788-7

8. Dalton, E., Griffin, D., Gallagher, T. F., de Vetten, N., & Milbourne, D. (2013). The effect of pyramiding two potato cyst nematode resistance loci to Globodera pallida Pa2/3 in potato. Molecular Breeding, 31(4), 921–930. 10.1007/s11032-013-9845-9

9. Den Nijs, L. (2024). NVWA-protocol: Uitvoeringsprotocol resistentieonderzoek aardappelmoeheid. https://www.nvwa.nl/documenten/plant/plantziekte-en-plaag/plantziekte/aardappelmoeheid/uitvoeringsprotocol-aardappelmoeheid-resistentietoetsing#:∼:text=In%20dit%20uitvoeringsprotocol%20vindt%20u,pallida%20moeten%20voldoen.

10. Den Nijs, L., & Van Heese, E. (2019). Development of new virulent populations in The Netherlands: How to detect this phenomenon in potato cyst nematodes (PCN)? Aspects of Applied Biology, 142. https://www.cabidigitallibrary.org/doi/epdf/10.5555/20220386633

11. Deng, L., Hao, J., Feng, J., Zhao, J., Dao, J., Xu, G., Zhou, K., Xu, Y., Wang, W., Zhang, S., Liu, C., Chen, M., Yang, Y., & Liu, X. (2025). Transcriptomic analysis of Phytophthora infestans races and evaluation of their pathogenicity on potato. BMC Plant Biology, 25(1), 716. 10.1186/s12870-025-06736-y

12. Ellenby, C. (1952). Resistance to the Potato Root Eelworm, Heterodera rostochiensis Wollenweber. Nature, 170(4337), 1016. 10.1038/1701016a0

13. Enciso-Maldonado, G. A., Lozoya-Saldaña, H., Colinas-Leon, M. T., Cuevas-Sanchez, J. A., Sanabria- Velázquez, A. D., Bamberg, J., & Raman, K. V. (2022). Assessment of Wild Solanum Species for Resistance to Phytophthora infestans (Mont.) de Bary in the Toluca Valley, Mexico. American Journal of Potato Research, 99(1), 25–39. 10.1007/s12230-021-09856-x

14. Eoche-Bosy, D., Gauthier, J., Juhel, A. S., Esquibet, M., Fournet, S., Grenier, E., & Montarry, J. (2017). Experimentally evolved populations of the potato cyst nematode Globodera pallida allow the targeting of genomic footprints of selection due to host adaptation. Plant Pathology, 66(6), 1022–1030. 10.1111/ppa.12646

15. EPPO. (2013). PM 7/119 (1) Nematode extraction. EPPO Bulletin, 43(3), 471–495. 10.1111/epp.12077

16. Ernst, K., Kumar, A., Kriseleit, D., Kloos, D. U., Phillips, M. S., & Ganal, M. W. (2002). The broad-spectrum potato cyst nematode resistance gene (Hero) from tomato is the only member of a large gene family of NBS-LRR genes with an unusual amino acid repeat in the LRR region. Plant Journal, 31(2), 127–136. 10.1046/j.1365-313X.2002.01341.x

17. Finkers-Tomczak, A., Danan, S., Van Dijk, T., Beyene, A., Bouwman, L., Overmars, H., Van Eck, H., Goverse, A., Bakker, J., & Bakker, E. (2009). A high-resolution map of the Grp1 locus on chromosome v of potato harbouring broad-spectrum resistance to the cyst nematode species Globodera pallida and Globodera rostochiensis. Theoretical and Applied Genetics, 119(1), 165–173. 10.1007/s00122-009-1026-1

18. Fournet, S. (2022, May). 30 years of research on resistance against Globodera pallida: an overview. ICN. https://www.youtube.com/watch?v=C3ZLULyyBmU&t=2279s

19. Fournet, S., Kerlan, M. C., Renault, L., Dantec, J. P., Rouaux, C., & Montarry, J. (2013). Selection of nematodes by resistant plants has implications for local adaptation and cross-virulence. Plant Pathology, 62(1), 184–193. 10.1111/j.1365-3059.2012.02617.x

20. Garrison, E., & Marth, G. (2016). Haplotype-based variant detection from short-read sequencing.

21. Gartner, U., Hein, I., Brown, L. H., Chen, X., Mantelin, S., Sharma, S. K., Dandurand, L. M., Kuhl, J. C., Jones, J. T., Bryan, G. J., & Blok, V. C. (2021). Resisting Potato Cyst Nematodes With Resistance. In Frontiers in Plant Science (Vol. 12). Frontiers Media S.A. 10.3389/fpls.2021.661194

22. Gebhardt, C., & Valkonen, J. P. T. (2025). ORGANIZATION OF GENES CONTROLLING DISEASE RESISTANCE IN THE POTATO GENOME. www.annualreviews.org.

23. Ghislain, M., Byarugaba, A. A., Magembe, E., Njoroge, A., Rivera, C., Román, M. L., Tovar, J. C., Gamboa, S., Forbes, G. A., Kreuze, J. F., Barekye, A., & Kiggundu, A. (2019). Stacking three late blight resistance genes from wild species directly into African highland potato varieties confers complete field resistance to local blight races. Plant Biotechnology Journal, 17(6), 1119–1129. 10.1111/pbi.13042

24. Jacobs, M. M. J., Vosman, B., Vleeshouwers, V. G. A. A., Visser, R. G. F., Henken, B., & van den Berg, R. G. (2010). A novel approach to locate Phytophthora infestans resistance genes on the potato genetic map. Theoretical and Applied Genetics, 120(4), 785–796. 10.1007/s00122-009-1199-7

25. Jones, F. G. W., Carpenter, J. M., Parrott, D. M., Stone, A. R., & Trudgill, D. L. (1970). Potato Cyst Nematode: One Species or Two ? Nature, 227(5253), 83–84. 10.1038/227083a0

26. Jones, J. T., Haegeman, A., Danchin, E. G. J., Gaur, H. S., Helder, J., Jones, M. G. K., Kikuchi, T., Manzanilla- López, R., Palomares-Rius, J. E., Wesemael, W. M. L., & Perry, R. N. (2013). Top 10 plant-parasitic nematodes in molecular plant pathology. Molecular Plant Pathology, 14(9), 946–961. 10.1111/mpp.12057

27. Kerlan, M.-C., Dantec, J.-P., Renault, L., Queré, J., Dantec, M.-A., Caromel, B., Montarry, J., & Fournet, S. (2017, July). Which association of QTL to obtain durable resistance to the nematode Globodera pallida? 20th EAPR Triennial Conference.

28. Kokot, M., Dlugosz, M., & Deorowicz, S. (2017). KMC 3: counting and manipulating k-mer statistics. Bioinformatics (Oxford, England), 33(17), 2759–2761. 10.1093/bioinformatics/btx304

29. Kreike, C. M., De Koning, J. R. A., Vinke, J. H., Van Ooijen, J. W., Gebhardt, C., & Stiekema, W. J. (1993). Mapping of loci involved in quantitatively inherited resistance to the potato cyst-nematode Globodera rostochiensis pathotype Rol. In Theor Appl Genet (Vol. 87). Springer-Verlag.

30. Leuenberger, J., Esnault, F., Lebas, P. L., Fournet, S., Cann, M. P., Marhadour, S., Prodhomme, C., Pilet-Nayel, M. L., & Kerlan, M. C. (2025). Identification by GWAS of marker haplotypes relevant to breed potato for Globodera pallida resistance. Theoretical and Applied Genetics, 138(3), 52. 10.1007/s00122-024-04794-8

31. Li, H., & Durbin, R. (2009). Fast and accurate short read alignment with Burrows-Wheeler transform. Bioinformatics, 25(14), 1754–1760. 10.1093/bioinformatics/btp324

32. Mundt, C. C. (2018). Pyramiding for Resistance Durability: Theory and Practice. Phytopathology®, 108(7), 792– 802. 10.1094/PHYTO-12-17-0426-RVW

33. Murashige, T., & Skoog, F. (1962). A revised medium for rapid growth and bio assays with tobacco tissue cultures. Physiologia Plantarum, 15(3).

34. Niere, B., Krüssel, S., & Osmers, K. (2014). Auftreten einer außergewöhnlich virulenten population der Kartoffelzystennematoden. Journal Für Kulturpflanzen, 55, 426–427.

35. Oerke, E. C. (2006). Crop losses to pests. In Journal of Agricultural Science (Vol. 144, Number 1, pp. 31–43). 10.1017/S0021859605005708

36. Paal, J., Henselewski, H., Muth, J., Meksem, K., Menéndez, C. M., Salamini, F., Ballvora, A., & Gebhardt, C. (2004). Molecular cloning of the potato Gro1-4 gene conferring resistance to pathotype Ro1 of the root cyst nematode Globodera rostochiensis, based on a candidate gene approach. Plant Journal, 38(2), 285–297. 10.1111/j.1365-313X.2004.02047.x

37. Pham, G. M., Hamilton, J. P., Wood, J. C., Burke, J. T., Zhao, H., Vaillancourt, B., Ou, S., Jiang, J., & Buell, C. R. (2020). Construction of a chromosome-scale long-read reference genome assembly for potato. GigaScience, 9(9), giaa100. 10.1093/gigascience/giaa100

38. Price, J., Preedy, K., Young, V., Todd, D., & Blok, V. C. (2024). Stacking host resistance genes to control Globodera pallida populations with different virulence. European Journal of Plant Pathology, 168(2), 373–381. 10.1007/s10658-023-02761-5

39. Prodhomme, C., Esselink, D., Borm, T., Visser, R. G. F., van Eck, H. J., & Vossen, J. H. (2019). Comparative Subsequence Sets Analysis (CoSSA) is a robust approach to identify haplotype specific SNPs; mapping and pedigree analysis of a potato wart disease resistance gene Sen3. Plant Methods, 15(1), 60. 10.1186/s13007-019-0445-5

40. Rigney, B., Blok, V., Griffin, D., Dalton, E., & Milbourne, D. (2017). Consistent action of two partially effective loci conferring resistance to Globodera pallida Pa2/3 across multiple nematode field populations. Plant Pathology, 66(6), 1031–1040. 10.1111/ppa.12640

41. Rouppe van der Voort, J., van der Vossen, E., Bakker, E., Overmars, H., van Zandvoort, P., Hutten, R., Klein Lankhorst, R., & Bakker, J. (2000). Two additive QTLs conferring broad-spectrum resistance in potato to Globodera pallida are localized on resistance gene clusters. Theoretical and Applied Genetics, 101, 1122–1130. https://link-springer-com.ezproxy.library.wur.nl/article/10.1007/s001220051588

42. Schaveling, A. S., te Molder, D. M., Heeres, P., van Steenbrugge, J. J. M., van de Ruitenbeek, S. J. S., van Schaik, C. C., van den Elsen, S., Smant, G., & Sterken, M. G. (2026). The potato cyst nematode Globodera pallida overcomes major potato resistance through selection on standing variation at a single locus. New Phytologist. 10.1111/nph.70886

43. Schaveling, A. S., van Rijt, L., Do, Y., Soffree, N., Langendoen, D., Room, H., Machado Bertran, A., Raven, M., van Kessel, S. P., van Heese, E. Y. J., van de Ruitenbeek, S. J. S., van Schaik, C. C., Kiewnick, S., Smant, G., & Sterken, M. G. (2026). Globodera pallida virulence on major potato resistance has a common genetic basis across Western Europe. PLOS Pathogens, 22(5 May). 10.1371/journal.ppat.1014201

44. Shanmugavel, S., Hu, S., Yang, H., Chen, S., Bamberg, J., Zhao, D., Basnet, B. R., Beil, C. T., Sheehan, M., Sathuvalli, V., & Wang, X. (2026). A chromosome-level genome assembly of Solanum brevicaule (PI 473011) Y1- 5, a wild potato relative with robust resistance to potato cyst nematodes. The Plant Genome, 19(2), e70265. 10.1002/tpg2.70265

45. Sikora, R. A., Molendijk, L. P. G., & Desaeger, J. (2021). Integrated nematode management and crop health: Future challenges and opportunities. In Integrated nematode management: state-of-the-art and visions for the future (1st ed., Vol. 1, pp. 3–10). CABI. https://cabidigitallibrary.org

46. Steiner, A. A. (1961). A universal method for preparing nutrient solutions of a certain desired composition. Plant and Soil, 15(2). 10.1007/BF01347224

47. Tammes JE, & Langendoen D. (2025). Resistentieveredeling tegen aardappelmoeheid bij Averis. Gewasbescherming, 56(6), 202–203.

48. Tan, M. Y. A., Park, T.-H., Alles, R., Hutten, R. C. B., Visser, R. G. F., & van Eck, H. J. (2009). GpaXItarloriginating from Solanum tarijense is a major resistance locus to Globodera pallida and is localised on chromosome 11 of potato. Theoretical and Applied Genetics, 119(8), 1477–1487. 10.1007/s00122-009-1149-4

49. Tarasov, A., Vilella, A. J., Cuppen, E., Nijman, I. J., & Prins, P. (2015). Sambamba: fast processing of NGS alignment formats. Bioinformatics, (31), 2032–2034. 10.5281/zenodo.13200

50. Trudgill, D. L. (1985). Potato cyst nematodes: a critical review of the current pathotyping scheme. Bulletin OEPP EPPO Bulletin, 15, 273–279.

51. Turner, S. J., Fleming, C. C., Moreland, B. P., & Martin, T. J. G. (2009). Variation in hatch among pathotypes of the potato cyst nematodes, globodera rostochiensis and G. pallida, in response to potato root diffusate from solanum spp. I. preliminary assessments to establish optimal testing conditions. Nematology, 11(5), 749–756. 10.1163/156854109X411005

52. Turner, S. J., & Rowe, J. A. (2006). Cyst Nematodes. In R. N. Perry & M. Moens (Eds.), Plant Nematology (1st ed., pp. 91–122). CABI. 10.1079/9781845930561.0091

53. Van Der Vossen, E. A. G., Van Der Voort, J. N. A. M. R., Kanyuka, K., Bendahmane, A., Sandbrink, H., Baulcombe, D. C., Bakker, J., Stiekema, W. J., & Klein-Lankhorst, R. M. (2000). Homologues of a single resistance-gene cluster in potato confer resistance to distinct pathogens: A virus and a nematode. Plant Journal, 23(5), 567–576. 10.1046/j.1365-313X.2000.00814.x

54. van Eck, H. J., Vos, P. G., Valkonen, J. P. T., Uitdewilligen, J. G. A. M. L., Lensing, H., de Vetten, N., & Visser, R. G. F. (2017). Graphical genotyping as a method to map Ny(o,n)stoand Gpa5 using a reference panel of tetraploid potato cultivars. Theoretical and Applied Genetics, 130(3), 515–528. 10.1007/s00122-016-2831-y

55. Varypatakis, K., Véronneau, P. Y., Thorpe, P., Cock, P. J. A., Lim, J. T. Y., Armstrong, M. R., Janakowski, S., Sobczak, M., Hein, I., Mimee, B., Jones, J. T., & Blok, V. C. (2020). The genomic impact of selection for virulence against resistance in the potato cyst nematode, globodera pallida. Genes, 11(12), 1–20. 10.3390/genes11121429

